# Temporal Vascular Endothelial Growth Factor Sub-type Gene Switching in SARS-CoV-related Inflammation - Basis for a Dual Gene Biomarker Approach

**DOI:** 10.1101/2022.11.06.515327

**Authors:** Asrar Rashid, Govind Benakati, Feras Al-Obeidat, Zainab A. Malik, Joe Brierley, Varun Sharma, Anuka Sharma, Love Gupta, Hoda Alkhazaimi, Guftar Shaikh, Ahmed Al-Dubai, Nasir Quraishi, Syed A. Zaki, Wael Hafez, Amir Hussain

## Abstract

This study examines temporal gene expression (GE) patterns in a murine model of SARS-CoV infection. We focused on a Temporal Gene Set (TGS) comprising pro-inflammatory genes (TNF, NFKB1, VEGF-A) and VEGF-B. A systematic search of the NCBI Geo database for MA15 (SARS-CoV) pulmonary studies using C57BL Wild (WT) mice and filtering according to TGS GE patterns eluded seven datasets for further analysis. Encompassing the GE profiles from these datasets alluded to a rising and falling pattern in TNF and NFKB1 GE. Also, our findings reveal a temporal decrease in VEGF-A GE coinciding with an increase in VEGF-B GE post-immunogenic stimulation. Notably, differential responses were observed with the MA15 dosage and in comparison, to other antigens (dORF6 and NSP16). Further, the human SARS-CoV-2 gene enrichment in this murine study confirms the MA15 murine model’s relevance for SARS research. Our study also suggests potential interactions between SARS-CoV-2 Spike protein and VEGF-related receptors, hinting at other pathophysiological mechanisms. Our results indicate severe inflammation may lead to a flattened VEGF-B GE response, influencing VEGF-B’s cell survival role. We underline the significance of considering VEGF-A/B interactions, particularly temporal differences, in manipulating angiogenic processes. Future research needs to consider temporal changes in VEGF-A and VEGF-B GE, in terms of time-associated gene-switching, in line with changing host inflammation.

## Introduction

Severe Acute Respiratory Syndrome (SARS) is caused primarily by coronaviruses, with the most notable, SARS-CoV-2, responsible for the Covid-19 pandemic. In adults, severe pulmonary disease, culminating in Acute Respiratory Distress Syndrome (ARDS), is a common outcome of SARS infection, particularly amongst high-risk groups with pre-existing conditions {Martínez-Colón, #4294}{Ravichandran, 2021 #4293}. Exploring mechanisms predisposing individuals to severe SARS, including the role of the virus’ spike protein and the Vascular Endothelial Growth Factor (VEGF), have been investigated {Harrison, 2020 #4295}{Yin, 2020 #3333}. Chi et al. detected higher VEGF levels in infected patients associated with the VEGF-A subclass {Chi, 2020 #3336}. Similarly, the MYSTIC study researching endothelial effects of Covid-19 infection demonstrated that blood VEGF-A levels were positively correlated with disease severity and ARDS development and also was the best discriminator in predicting 60-day in-hospital mortality {Rovas, 2021 #3332}. VEGF-A binding to VEGF receptors (VEGFR) increases vascular permeability, nitric oxide-mediated vasodilation, and angiogenesis {Stookey, 2020 #3354}. The angiogenic potential of VEGF-A is crucial in Covid-19, as demonstrated by studies revealing its role in the formation of luminal cylindrical microstructures and angiogenesis in pulmonary tissue {Ackermann, 2020 #3542}.

The role of other VEGF subtypes in SARS has not been well documented, including that of the VEGF-B subtype. The differential effects of VEGFR-1 and VEGFR-2 activation by VEGF-A and VEGF-B potentially contribute to their distinct roles in various physiological and pathological processes. Specifically, VEGF-A binding to VEGFR-1 does not significantly affect receptor activation; however, VEGF-B binding to VEGFR-1 promotes cell survival {Lal, 2018 #3622}. In pheochromocytoma studies, VEGF-B has been found to displace VEGF-A from VEGFR-1, indicating a shift of VEGF-A to VEGFR-2 {Abe, 2021 #4298}. This displacement effect may be the reason why VEGF-B binding to VEGFR-1 results in VEGF-A/vascular endothelial growth factor receptor (VEGFR) 2 (VEGFR2) pathway activation {Zafar, 2017 #4274}. VEGF-A binding to VEGFR-2 triggers cell migration, although receptor binding is of low affinity yet plays a vital role in angiogenesis transduction. It is worth noting that VEGFR-2 receptors are abundant in pulmonary tissue, underscoring the importance of VEGF-A in respiratory health and disease {Peach, 2018 #4273}. VEGF-B is co-expressed with VEGF-A in all tissues but is most abundant in tissues with high metabolic activity, such as brown adipose, heart, and skeletal muscle, the heart, and skeletal muscle {Olofsson, 1996 #4303}. These findings suggest that the effects of VEGF proteins on VEGFR-1 and VEGF-R2 receptors may contribute to pathogenesis.

VEGF-A exhibits pro-inflammatory effects important in the pathogenesis of severe SARS-CoV-2 infection. Inflammatory responses involving VEGF-A occur at three distinct points along the disease pathway {Dabravolski, 2022 #4300}. Initially, VEGF-A stimulates IL-6 activation, releasing STAT-3 and generating an autofeedback loop that amplifies VEGF-A production. Additionally, the Akt pathway supports feedback loop secretion via the IL-6/STAT-3 pathway. Finally, as a pro-inflammatory molecule, VEGF-A ultimately activates NF-κB. However, VEGF-B exhibits a strong affinity for VEGFR-1, effectively displacing VEGF-A upon receptor binding. Nevertheless, research on the function of VEGF-B in inflammatory diseases remains limited. A study of multiple sepsis datasets suggests an inverse association between VEGF-A and VEGF-B gene expression in sepsis (paper in review at ‘Shock Journal). Similar patterns have also been observed in Kawasaki Disease (unrelated to SARS-CoV-2 infection), highlighting changes in the transcriptome during inflammation based on gene expression of tumor necrosis factor (TNF) and NFKB1 (NFKB1){Rashid, 2022 #4475}. Given these studies and the ability of VEGF-A and VEGF-B to bind to VEGFR-1, these findings suggest the idea of temporal switching of VEGF-A and VEGF-B expression in SARS-associated inflammation.

Considering the global significance of SARS, there is a need to enhance disease biomarking for improved clinical precision. Therefore, this paper aims to investigate temporal gene expression patterns in a murine model of SARS-CoV infection using C57BL Wild (WT) mice. In this study, methodologies previously established in sepsis and Kawasaki Disease will be applied to analyze the SARS temporal transcriptome (references). Several publicly available transcriptomic datasets were utilized to explore the effects of nasal instillation of the MA15 strain of SARS-CoV in C57BL Wild (WT) mice. This MA15 strain of SARS-CoV is derived from the Urbani strain and was introduced into the murine model through serial passage via the respiratory tract of young BALB/c mice {Roberts, 2007 #3505}. The highly immunogenic nature of the MA15 strain is evidenced by its lethality after intranasal administration in the murine context. Intranasal instillation of MA15 in the murine respiratory tract leads to rapid viremia, high-titer viral replication in the lungs, and extrapulmonary spread. This is accompanied by neutropenia and lymphopenia, indicative of the severe nature of the viral infection. Insights obtained from this study on the temporal transcriptome in the murine MA15 pulmonary model using WT mice could contribute to a better understanding of human SARS-CoV pulmonary infection, particularly in terms of identifying biomarkers to track associated patterns of inflammation.

## Material and Methods

### Murine Dataset Selection

This research conducted temporal studies on murine pulmonary infection post-MA15 coronavirus exposure compared to controls. We focused on Murine WT studies which were temporal and related to the SARS disease model. Consequently, pertinent Gene Expression (GE) datasets were sourced from the NCBI GEO and EMBL-EBI databases. The experimental models in these studies utilized intranasal instillation of SARS MA15 Plaque Forming Units (PFU) in phosphate-buffered saline (PBS) versus Mock (PBS only). Mock samples were used as the control group to establish a baseline for comparison, allowing researchers to determine the effect of experimental variables. The analysis focused on C57BL/6 Murine Wildtype (WT) lung studies after MA15 SARS-CoV infection. The search strategy retrieved SARS-related gene expression data from Microarray experiments and RNA-seq datasets (Figure 1). Datasets with samples from more than one time point were included. The EMBL-EBI and NCBI GEO databases yielded 248 and 34 entries, respectively, leading to the selection of 13 microarray datasets. The emphasis was on comparing MA15 infection versus mock datasets but studies GSE49262 and GSE49263, as well as MA15, tested additional antigens. Datasets GSE68820 and GSE33266 yielded insights into temporal and antigen dosage effects on immune stimulation. The tested doses of MA15 PFU included 10^2, 10^3, 10^4, and 10^5. RNA-seq studies provided no qualifying datasets.

**Figure 1.**
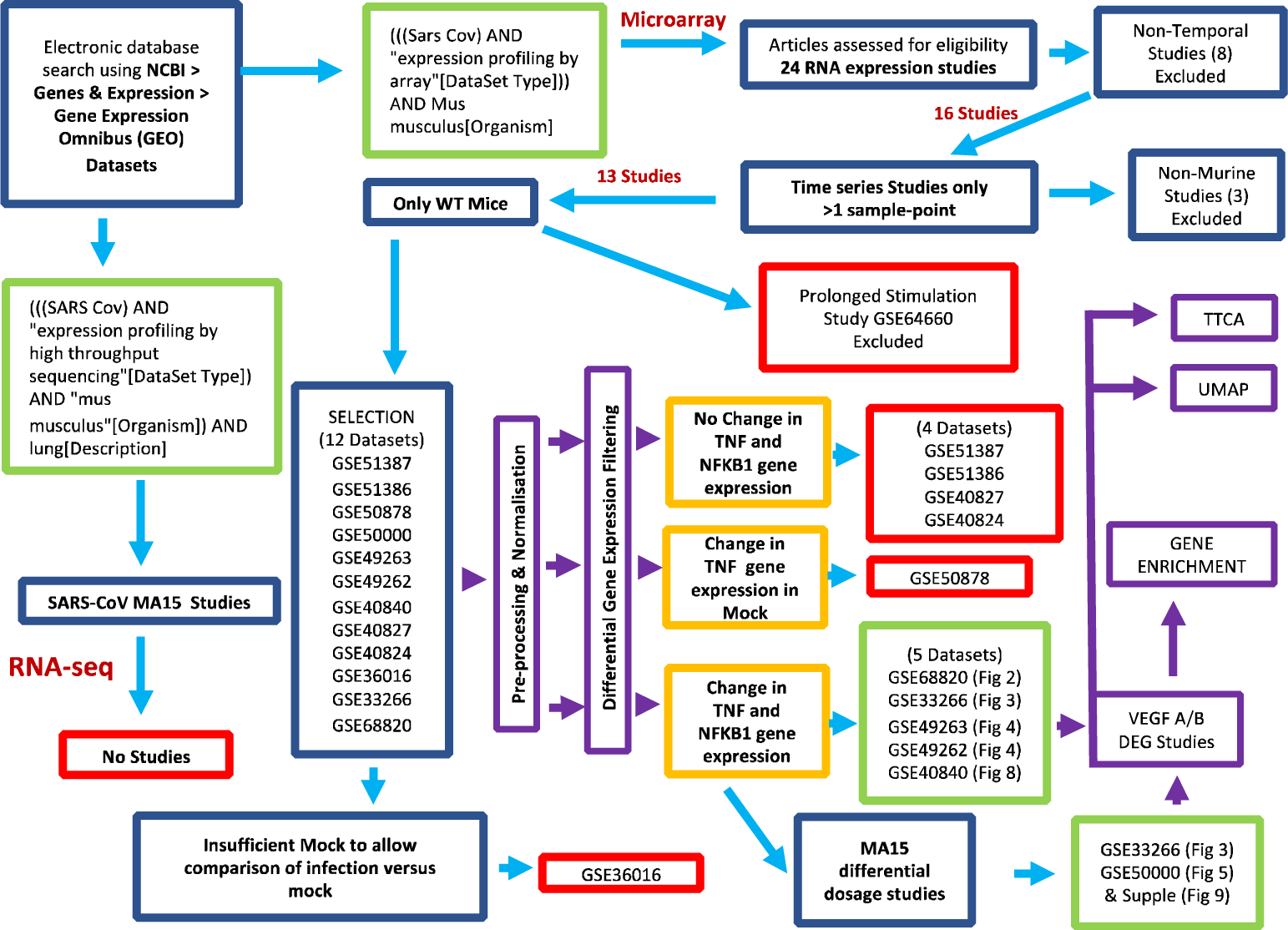
Selection of NCBI Datasets for Gene Expression Analysis in Murine MA15 Pulmonary Infection Studies, Including The Application of Methods. The search strategy undertaken on the 26^th^ of August 2022 used the NCBI GEO database and search criteria as shown. The goal was to identify temporal studies that use a one-event MA15 antigenic stimulation of the Murine respiratory tract. 24 items were found in which eight datasets researched non-temporal after Murine MA15 SARS CoV infection, thus were excluded from further analysis. Further, three of the sixteen datasets were unrelated to WT mice and were eliminated. Also excluded were dataset GSE64660 which was a prolonged MA15 exposure study, and GSE36016, in which there was insufficient mock to allow comparison against controls. Gene expression (GE) was classified according to TNF and NFKB1 categories; apart from the category where there was both TNF and NFKB1 differential GE, the other datasets were excluded. This led to 8 datasets being eligible for analysis, of which two datasets (GSE33266 and GSE50000) were dosage studies, testing different levels of MA15 dosing during stimulation. In purple are the Methods used in this paper, the first being pre-processing and normalization. This is followed by differential gene expression filtering based on TNF and NFKB1 gene expression. No change in TNF and NFKB1 GE is assumed to reflect that the transcript may (at the sample time) indicate not showing inflammation according to gene expression and was excluded from further examination. Or that Mock shows changes in TNF GE, suggesting not to be a true control, so that dataset was also removed excluded. Only datasets are indicative of TNF and NFKB1 GE changes are chosen for further analysis. As well as TNF and NFKB1 GE, the method includes an analysis of VEGF-A and VEGF-B differential GE (DGE) analysis. Datasets then undergo unsupervised analysis between infected samples and controls, resulting in a gene list, which undergoes gene enrichment and UMAP analyses.

### In Silico Statistical and Gene Ontology analyses

Qlucore Omics Explorer (QOE) version 3.7 was employed for Differential Expression of the Genes (DEGs) analysis, generating Principal Component Analysis (PCA) plots, and conducting comparisons and hierarchical clustering. Gene symbols were generated for probes, with expression data averaged on all microarray datasets for multiple results of the same Gene Symbol before box-plot analysis and GSEA. Genes were standardized, and the p-value and False Discovery Rate (FDR) ‘q’ were applied, with both values below 0.05 required for statistically significant. Hierarchical clustering in QOE was based on Euclidean distance and average linkage clustering. All microarray datasets were normalized and log2 transformed using R-script for further downstream analysis. Protein docking involved VEGF-A protein data bank structure receptor 1WDF and SARS-CoV spike protein-ligand 1BJ1 using the Barcelona supercomputing server pyDOC (PMID: 23661696). Over a hundred receptor and ligand variants were shortlisted for further analyses. Top hits from prediction models were plotted using Pymol{SchrödingerLLC, #4302}. Student’s t-test was employed to compare gene expression between sample groups, with p-value of <0.05 indicating statistical significance. Enrichment was performed via the EnrichR platform{Xie, 2021 #4350}, facilitating mapping onto key databases such as KEGG 2021. The platform also supported direct porting of plots to Appyter notebooks for various data visualizations, such as hexagonal plots from the KEGG_2021_Human gene set library. Each hexagon in the plot represents a single term, with similar gene sets grouped together. Brighter colors indicate higher Jaccard similarity, while significant overlap in the input query gene set manifests as blue hexagons in the database.

### Understanding dynamic changes in gene expression using Transcript Time Course Analysis and Box plot analysis

Two methods were employed to understand temporal dynamics in gene expression: t-tests for time point comparison and Transcript Time Course Analysis (TTCA) for temporal data visualization{Albrecht, 2017 #2534}. While t-tests evaluated changes in gene expression based on box plot analysis, TTCA offers a graphical depiction of gene expression shifts. Whereas box plots were applied to pre-assigned genes for temporal comparison, TTCA is an unsupervised approach. Over-representation analyses, utilizing hypergeometric distribution-based testing, were conducted on TTCA-derived outcomes. Following this, gene expression for TNF, NFKB1, VEGFA, and VEGFB was curated using the Hugo database. This facilitated the representation of significant genes under categories such as ‘Consensus,’ ‘Early Response,’ ‘Middle Response,’ ‘Late Response,’ ‘Complete Response,’ ‘Dynamic,’ and ‘MaxDist.’

## Results

Thirteen Murine datasets (with >1 temporal sampling point) studying pulmonary infection by MA15 in the WT Murine model were selected according to the search strategy shown (Figure 1).

### A. Murine SARS Gene Expression Analysis Dataset GSE68820

The dataset GSE68820 included an analysis of 27 WT samples, as depicted in Figure 2. Temporal Differential Gene Expression (DGE) analysis comparing Day 2 Days Post Instillation (DPI) with 7 DPI revealed a significant decrease in TNF (p=1.858e-05) and VEGF-A (p=7.617e-03), accompanied by an increase in VEGF-B gene expression (p=7.580e-07) (Figure 2A). NFKB1 did not show any significant DGE (p=1.387e-01). The temporal trends from 1 to 7 DPI of MA15 suggested an inverse association between VEGF-A and VEGF-B gene expression. TNF and VEGF-A gene expression demonstrated a downward trend over the study period, indicating a decrease in VEGF-A expression as inflammation subsided, with an opposite increase trend in VEGF-B DGE.

**Figure 2.**
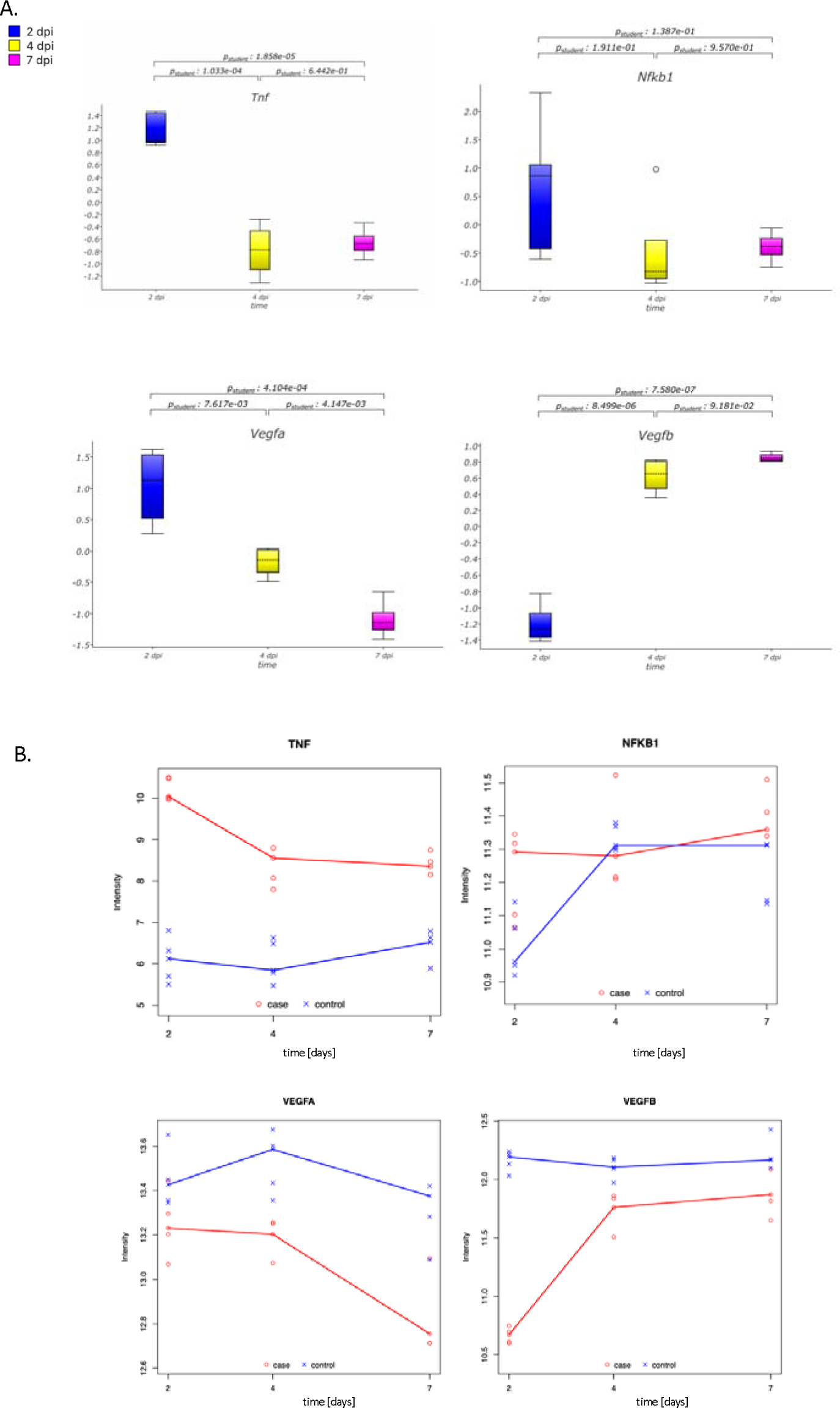
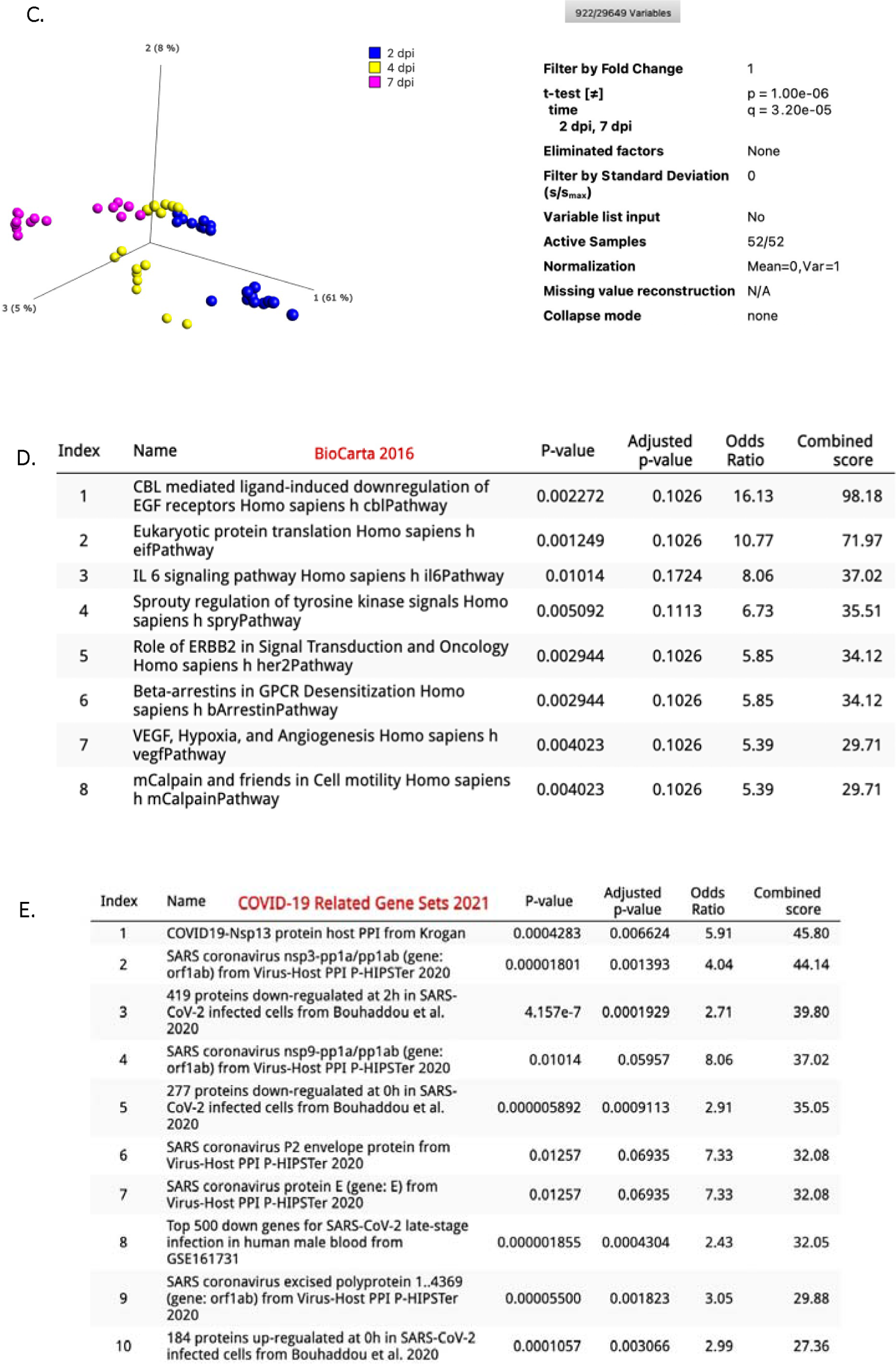
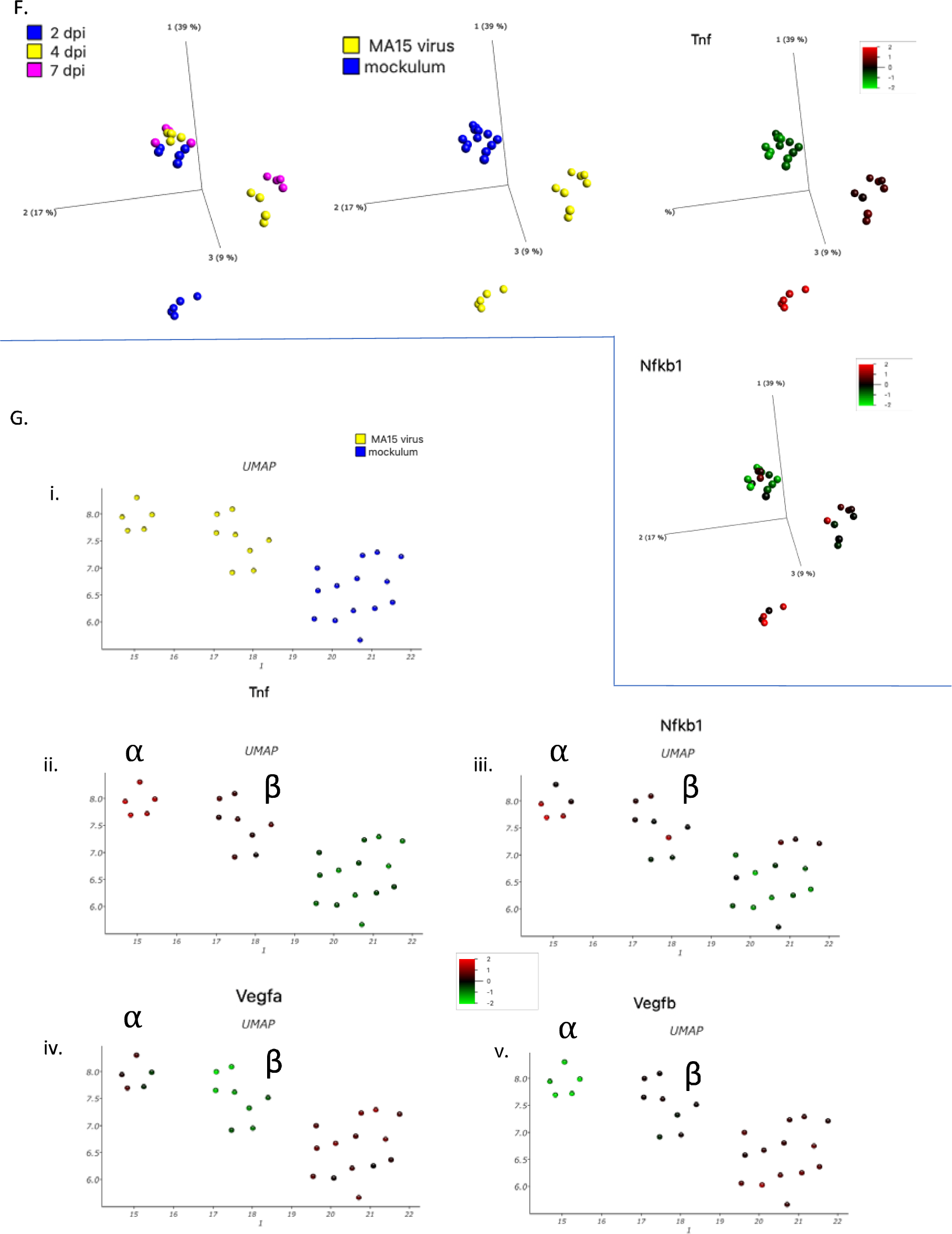
A study of Pulmonary Gene Expression in Murine (WT) Model with CoV virus nasal instillation of MA15 versus Mock. This figure illustrates the results of an analysis conducted on murine (WT) lung tissue (GSE68820) from a pulmonary study. 27 WT samples were assessed within the GSE68820 dataset and averaged to obtain 15,961 variables. The comparison was drawn between MA15 and Mock nasal instillation. Notably, on days 2 and 7, VEGF-A, TNF, and NFKB1 gene expression was significantly down-regulated, whereas VEGF-B was markedly up-regulated in the group suffering from MA15 pulmonary infection (Figure 2A). Trends in VEGFA and VEGFB gene expression are suggested by TTCA plots (Figure 2B). TTCA also highlights an increased intensity for TNF gene expression (from day two to day seven) compared to controls, whereas NFKB1 gene expression appears to converge between controls and cases by day four. A comparison between two and seven Days Post Instillation (DPI) resulted in a t-test on the 52 samples, filtering to 922 genes (p=1.00e-06 and q= 3.20e-05), featuring QOE in a non-collapsed mode (Figure 2C). The parsed gene list through the Enrichr online platform (https://maayanlab.cloud/Enrichr/enrich<u>#</u>) identified 720 unique genes. Pathway enrichment analysis revealed BioCarta2016 pathways that include IL-6 signaling and ‘VEGF, Hypoxia and Angiogenesis’ (Figure 2D). Moreover, Enrichr highlighted pathways containing Covid-19 related genes (Figure 2E). A PCA representation of the genes is depicted (Figure 2F). Subsequently, a 2D UMAP plot was performed using the averaged dataset (Mean=0, Var =1, Perplexity =11, with default settings, number of neighbors = 15, minimum distance =0.1, SVD dimensions = 50, and Random State = 42) (Figure 2G). This unsupervised method led to the formation of the clusters observed, encompassing both the mock and infected groups (Alpha and Beta). The Alpha MA15 stimulated group displayed a clear distinction between TNF up-regulation and VEGF-B down-regulation, in contrast to the Beta group, which exhibited VEGF-B down-regulation.

Furthermore, the changing gene expression dynamics were examined using Time-series Transcriptome Cluster Analysis (TTCA), comparing MA15 cohorts against mock controls (Figure 2B). TNF intensity patterns displayed differences between MA15 cases and controls, while initial NFKB1 gene expression (GE) showed separate patterns between Mock and controls, eventually converging by 4 DPI. The gene expression intensities of VEGF-A and VEGF-B remained divergent between cases and controls. However, over the study period, the intensity of VEGF-A decreased while the intensity of VEGF-B increased, approaching levels observed in the Mock (control) group.

A t-test comparison between 2 and 7 DPI (p=1.00e-06 and q= 3.20e-05) resulted in the identification of 922 genes showing differential change. To further analyze these genes, gene enrichment analysis was conducted and mapped onto the BioCarta2016 database (Figure 2D), revealing enrichment in the IL-6 signaling pathway. The gene list was also mapped onto the Covid-19 Related Gene Sets 2021, demonstrating the enrichment of several SARS-related pathways (Figure 2E). These findings align with human Covid-19 (clinical) studies, where elevated IL-6 levels have been associated with poor clinical outcomes {Coomes, 2020 #4351}. Furthermore, the enrichment analysis of the 922 genes highlighted the presence of the SARS coronavirus P2 envelope protein, suggesting a potential overlap in pathogenesis between the SARS-CoV MA15 virus in mice and SARS-CoV-2 in humans.

In the case of GSE68820, a 2D UMAP (Uniform Manifold Approximation and Projection) plot was generated (Figure 2G). The UMAP plot effectively distinguished MA15 samples from mock samples, with MA15 samples further divided into two distinct categories (Figure 2Gi-iv). The alpha grouping within the MA15 samples displayed up-regulation of TNF and downregulation of VEGF-B. In contrast, the mock samples exhibited downregulation of TNF, and the other gene expression patterns were inconclusive.

### B. Temporal Murine SARS Gene Expression Analysis Studies study (GSE33266)

The impact of MA15 on the pulmonary bed was investigated using a temporal MA15 dose stimulation dataset (GSE33266, Figure 3). At the lowest administered dose of MA15 (10^2^ PFU), GE increased for TNF, and for NFKB1 (1 DPI and 4PI), there was an increase, followed by a decrease in GE (4 and 7 DPI). However, as the MA15 dose escalated from 10^3^ to 10^4^ PFU, peak gene expression for TNF and NFKB1 was observed at 1-day post-infection (DPI). Interestingly, no differential GE for TNF or NFKB1 was recorded at the maximum dose of 10^5^ PFU. In the case of VEGF-A, a consistent decrease in GE was observed starting from a dose of 10^3^ PFU (1 DPI versus 7 DPI). VEGF-B GE showed a falling trend (1 and 7 DPI; 2 and 4 DPI) at 10^3^ PFU. However, at an escalated dose range (10^4^ PFU), the GE of VEGF-B initially exhibited a decline, followed by an increase in GE resembling a ‘V’ shape. A similar ‘V’ pattern in the VEGF-B GE profile was evident at 10^5^ PFU.

**Figure 3:**
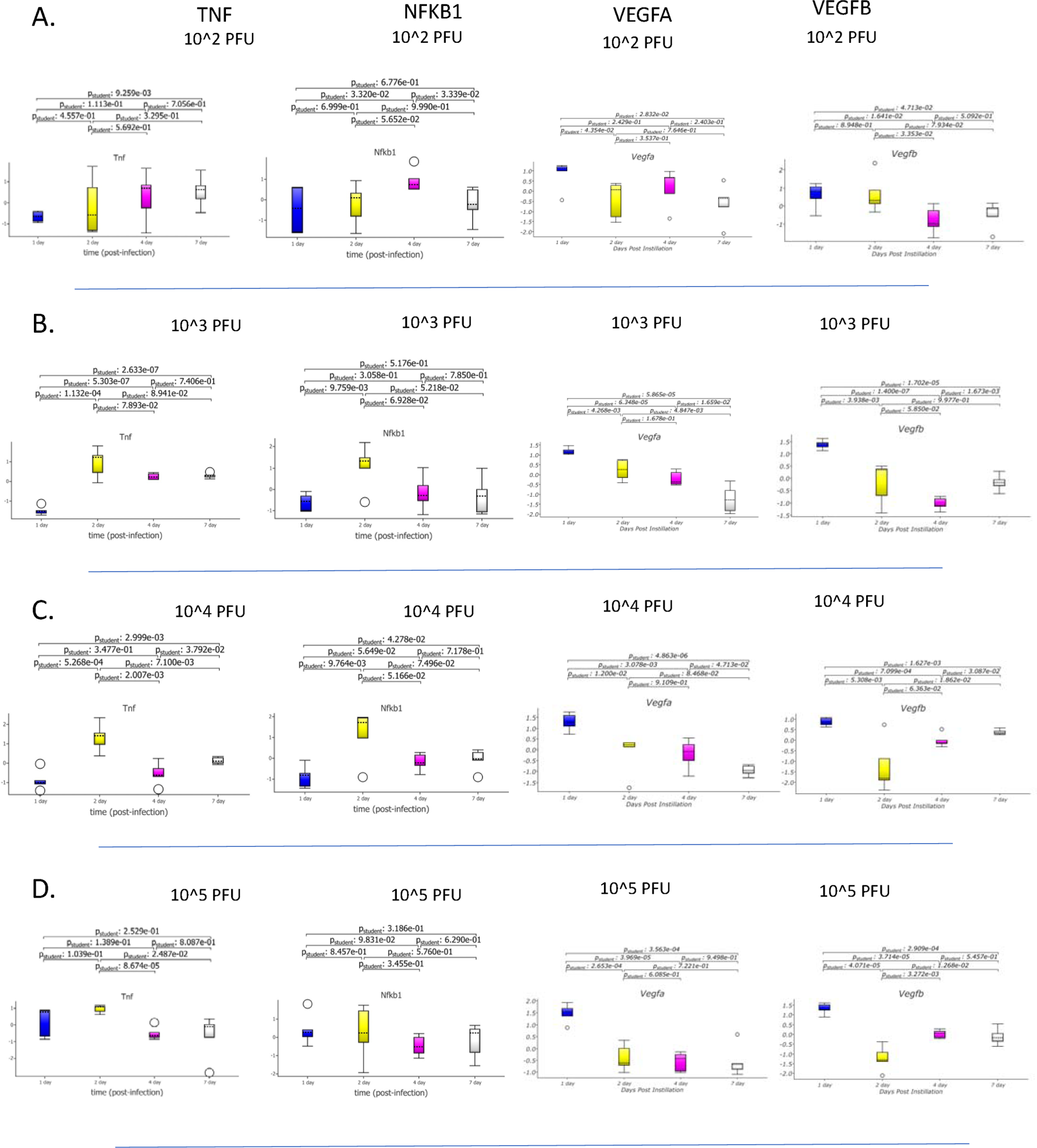
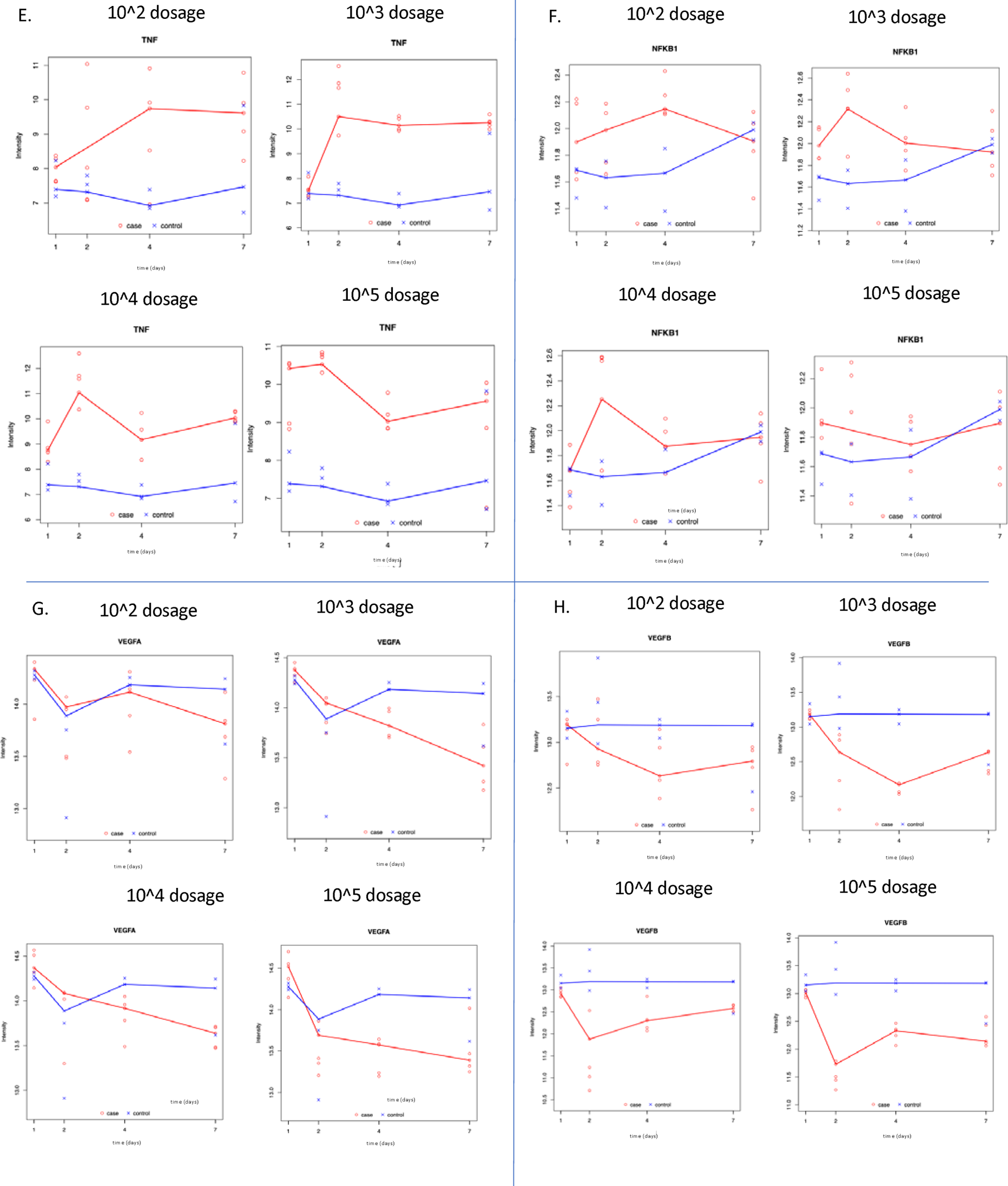
Murine Pulmonary effects of Varying MA15 doses as assessed by VEGF-A, VEGF-B, TNF, and NFKB1 gene expression. **A-D**. A temporal study (**GSE33266)** is shown after SARS COV MA15 nasal instillation at varying MA15 doses (10^2, 10^3, 10^4, and 105). NFKB1, TNF, VEGFA, and VEGFB gene expression (GE) are illustrated by TTCA (**A-D**) and box plot (**E-H**) analysis. TTCA **VEGF-A** GE diverges in intensity between controls (Mock) and cases (MA15 stimulated) in intensity (Day 4 to 7). TTCA **VEGF-B** GE: TTCA **TNF** GE showed a divergence between cases (MA15) and controls (Mock), at all the dose levels remaining throughout the study period. At the lower doses (10^2 to 10^4), there is a divergence from 1 Day post-instillation, with the 10^5 dose showing the most initial divergence from Day 1. Whereas, for **TTCA/NFKB1** GE, controls and cases converge with increasing time. Trends in NFKB1 and TNF gene expression showed 10^3 and 10^4 PFU dosing, resulting in an increase in NFKB1 gene expression between day 1 and day 2, suggesting an acute inflammatory response. Box plot **VEGF-A** gene expression: 10^2 dose (no change GE), at 10^3^, 10^4^, and 10^5^, a significant decrease in GE is noted (Day 1 compared to Day 7). Box plot **VEGF-B** gene expression: at 10^2 (no change GE), for 10^3^, a significant fall is noted at 10^3^, 10^4^, and 10^5^doses (Day 1 to Day 7); however, both the 10^4^ and 10^5^ doses show a ‘v’ shape at Day 1, Day 2 and Day 4 vertices.

The TTCA analysis revealed that NFKB1 gene expression initially exhibited an increase in cases (MA15) compared to controls (Mock) (Figure 3). TTCA patterns for TNF gene expression displayed intensity differences between cases (MA15 stimulated group) and controls. In the case of TNF, by the end of the study period, TNF GE remained significantly high in cases compared to controls. For NFKB1, there was an initial disparity between cases and controls, but by day 7 (in contrast to TNF GE), the intensities converged between infection and mock. For the 10^2, 10^3, and 10^4 dose, TTCA suggested a peak in GE for TNF and NFKB1 between 1 DPI and 2 DPI. This was consistent with DGE from 1DPI and 2DPI by box plot (at 10^3 and 10^4 PFU).

In TTCA, VEGF-A GE diverged between cases and controls, with decreasing GE intensities observed in the cases across time. As for VEGF-B, there was an initial divergence between cases and controls as VEGF-B intensities declined, followed by a convergence of the two groups (10^2, 10^3, and 10^4 dose).

### C. MA15 versus Mock Pulmonary Gene Expression GSE40840

GSE40840 data analysis allowed a comparison of MA15 versus Mock stimulation in the WT murine model (Figure 8). Comparing gene expression at 4DPI (Days Post-Infection) versus 7DPI, significant up-regulation was observed for TNF (p=4.898e-02) and NFKB1 (p=3.83e-06), VEGF-A showed downregulation in GE (p=1.484e-03) with no change in VEGF-B GE (p=8.545e-01).

### D. Antigen Variation on MURINE SARS Gene Expression Analysis GSE49262 and GSE49263

The effects of MA15 on gene expression (GE) were compared to other antigens (dORF6:GSE49262 and NSP16GSE49263) (Figures 4A and B). The Mock samples showed no significant differential GE between 1 DPI and 7 DPI in both datasets. For both datasets, dORF6, NSP16, and MA15 induced a decrease in TNF, NFKB1, and VEGF-A GE. Furthermore, compared to dORF6, MA15 stimulation caused greater downregulation in TNF and VEGF-A GE for both datasets. According to the p-values, the largest decrease in GE magnitude was observed between day 1 and day 4 for MA15 compared to dORF6 and NSP16 for all four genes analyzed across similar time periods in both datasets. Regarding VEGF-B, only dORF6 (in GSE49262) resulted in a v shape change in VEGF-B GE with an inflection at 2 DPI. Whereas MA15 resulted in no differential gene expression response observed for VEGF-B GE.

**Figure 4:**
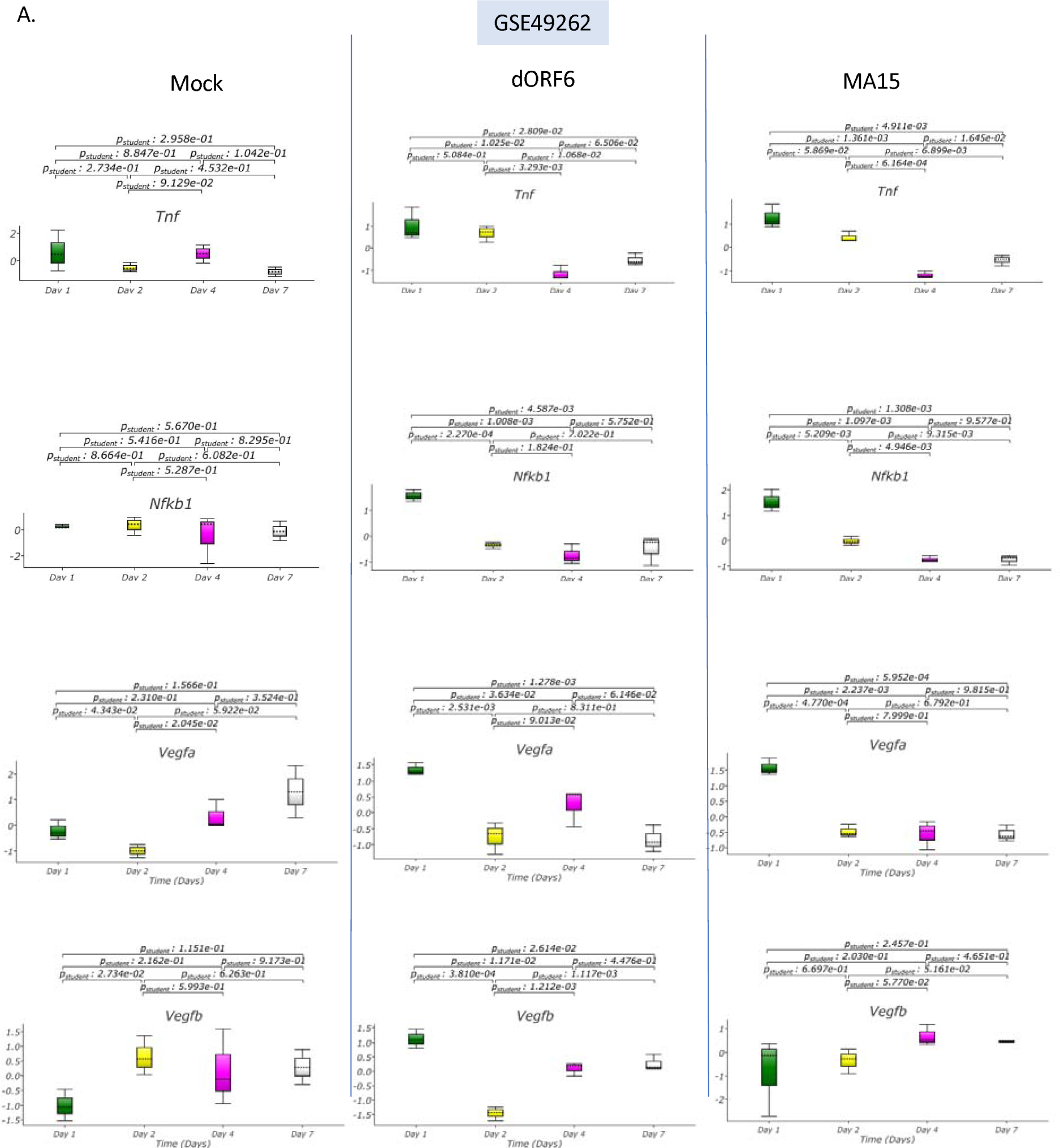
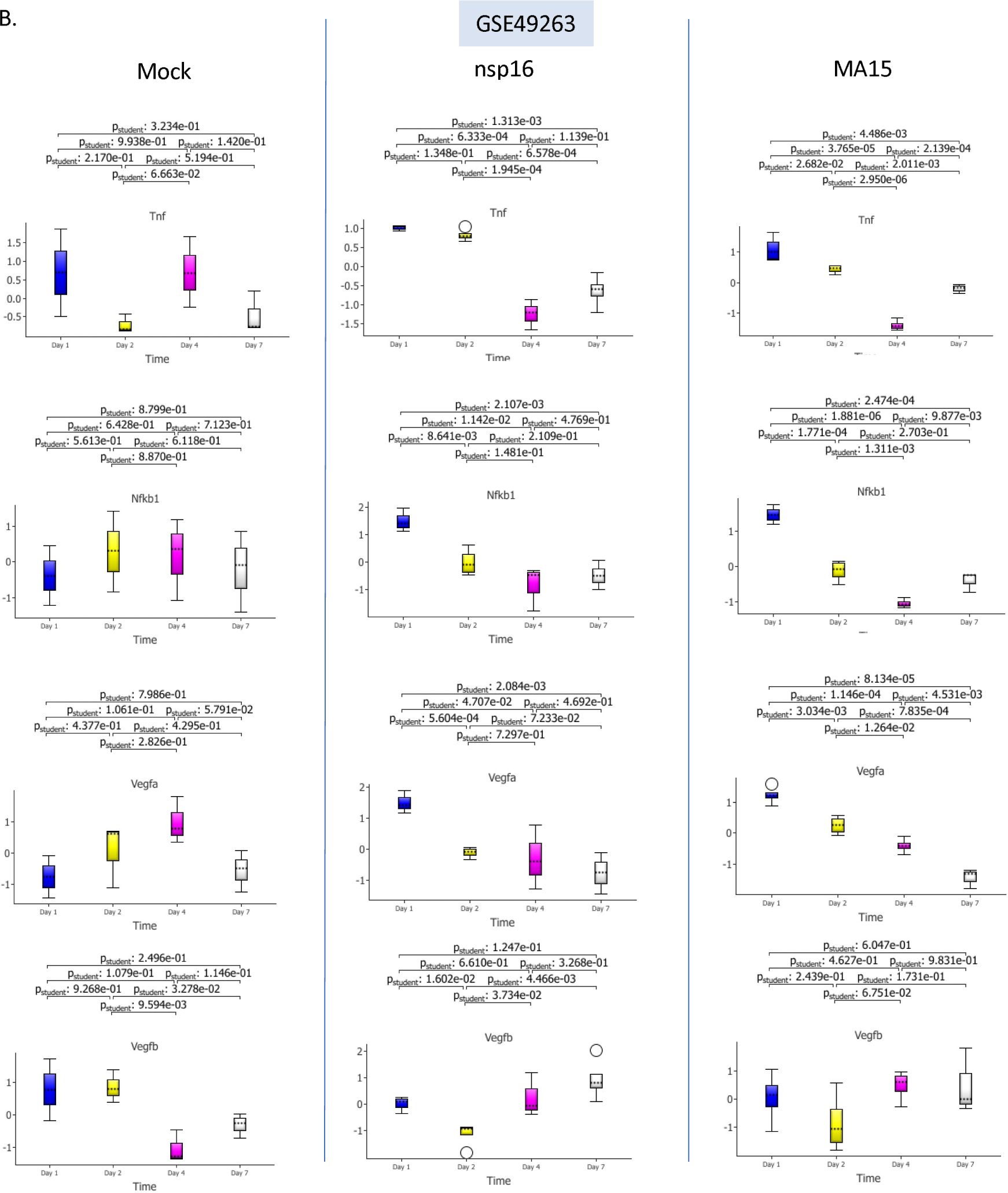
Effect of Differing Antigenic Patterns of Pulmonary Gene Expression in the WT Murine Model. Stimulation of WT Mice with either Mock, dORF6, or MA15 is shown in GSE49262 (Figure 4A) and GSE49263 (Figure 4B). For each of the genes (TNF, NFKB1, VEGF-A, and VEGF-B), Mock samples showed no change in GE at the start and end of the study. **Dataset GSE49262** showed TNF and NFKB1 GE to fall from day 1 to day 4 for dORF6 and MA15. On Day 1 compared to Day 7, there is a significant fall in VEGF-A GE for both dORF6 and MA15 GE. Also, for VEGF-B, only dORF6 shows a significant fall between Day 1 and Day 2, followed by a significant rise in GE between Day 2 and 7. For MA15, there is no change in VEGFB GE. For **Dataset GSE49263,** Mock Samples show no significant change in GE for each gene (TNF, NFKB1, VEGF-A, and VEGF-B). For NSP16 and MA15, GE for TNF, NFKB1, and VEGFA shows a significant fall when comparing Day 1 and Day 4 samples. For NSP16 and MA15, there is no significant change in GE between the start and end of the study.

### E. Analysis of changes in TNF, NFKB1, VEGF-A, and VEGF-B in dose and antigen studies (GSE50000)

UMAP analysis on the dataset GSE50000 was performed, illustrating dose and antigen stimulation effects in the WT Pulmonary model (Figure 5). The UMAP diagram revealed the presence of three distinct clusters (Figure 5). Two clusters, alpha, and beta, were associated with MA15 samples, and an all-together cluster was related to Mock. Both the MA15 alpha and beta sub-groups exhibited an up-regulation of TNF and a downregulation of VEGF-B. However, for VEGF-A GE, the alpha cluster showed a down-regulation, while the beta cluster was up-regulated. Regarding NFKB1, the Beta cluster tended to show downregulation, while the direction of NFKB1 GE for the Alpha cluster was less clear.

**Figure 5:**
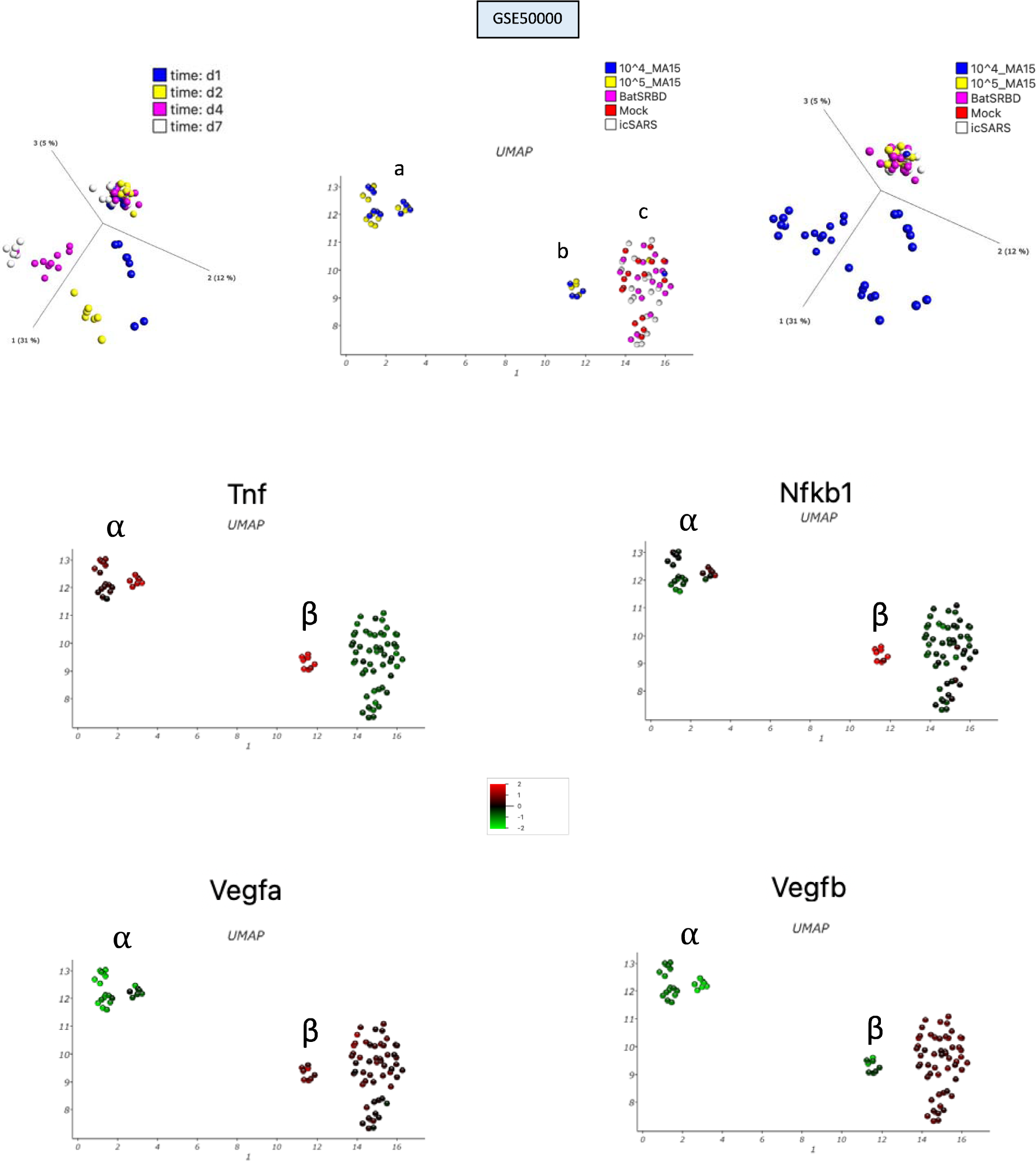
VEGFA, VEGFB Gene Expression, Pulmonary Murine WT model. For the GSE50000 dataset, a 2D UMAP plot was undertaken with the dataset averaged in QOE according to gene symbol (Mean=0, Var =1, Perplexity =11, with default settings, number of neighbors = 15, minimum distance =0.1, SVD dimensions = 50, and Random State = 42) (Figure 5A). This generated three groupings (**a**, **b**, and **c**) according to antigen type (collapsed).

Box plots illustrated GE profiles for the 10^4^ and 10^5^ MA15 doses (Figure 9). In both datasets, the pro-inflammatory genes NFKB1 and VEGF-A were down-regulated (1DPI compared to D7). However, there was TNF downregulation at the MA15 104 dose comparing study start and end points, but there was no significant change at the 10^5^ MA15 dose during the same time duration. However, this 10^5^ MA15 did induce a TNF GE effect with downregulation noted comparing GE between the 1DPI and 4DPI. At the 10^4^ MA15 dose for VEGF-B GE, there were no significant differences between the starting and ending points of the study, though a significant fall in VEGF-B GE was noted between 1 DPI to 2 DPI. Instead, for the 10^5^ MA15 dose, no significant changes in VEGF-B GE were observed.

### F. PROTEIN-PROTEIN DOCKING

A simulation exercise was performed to investigate the interaction between VEGF receptors and the spike protein of SARS-CoV (Figure 6). The pyDOC platform was utilized to predict the best docking models between VEGF receptors and the spike protein. These models were ranked based on factors such as electrostatics and desolvation energy (Table 3). All predicted models were extracted and plotted to assess the stability of the interactions. Remarkably, the top predicted models exhibited a stable interaction between VEGF receptors and the spike protein of SARS-CoV (Figures 6A-6J). showed downregulation in GE (p=1.484e-03).

**Figure 6.**
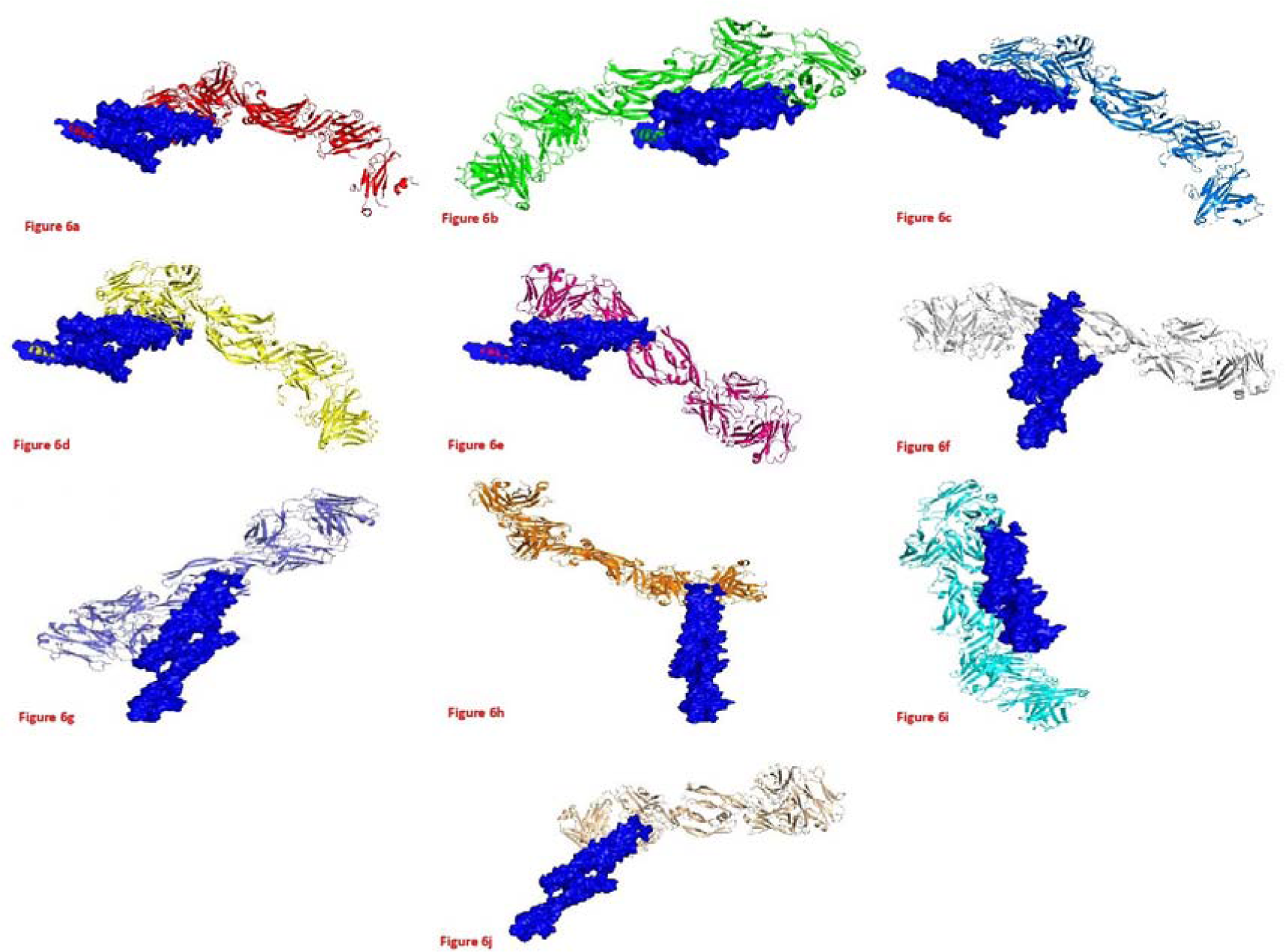
VEGF Receptors Docking for SARS-CoV Spike Proteins. Docking of VEGFA and VEGFB receptors were modeled and docked with ligands of Spike protein of SARS-CoV.

**TABLE 1:** Study characteristics of WT Pulmonary Study after MA15 nasal instillation. Ten and Twenty-week-old mice were infected by intranasal instillation of 10^5 or 10^4 PFU of SARS MA15 in 50 ¬µl of PBS or mock-infected with PBS alone. Lungs were harvested at the above time points according to Days Post Infection (DPI). The GSE number is the NCBI database identifier for the concerned study. GSE33266 and GSE68820 were studies performed by the same research group. GSE33266 concentrates on dose effects, and the GSE68820 on temporal changes, and their analysis data is shown (Figure 2 and Figure 3, respectively). In the study, *GSE36016 mock was only provided for day 5.

| GSE Number | Number of MA15 instilled animals | Number of Mock Instilled | The dose of SARS-CoV MA15 (PFU) Instilled | Age of Mice (weeks) | Time points (DPI) |
| --- | --- | --- | --- | --- | --- |
| 51387 | 5 | 4 | $10^5$ | 20 | 4,7 |
| 51386 | 7 | 8 | $10^4$ | 20 | 4,7 |
| 50878 | 8 | 9 | $10^5$ | 10 | 2,4,7 |
| 50000 | 32 | 16 | $10^4$ or $10^5$ | 20 | 1,2,4,7 |
| 49263 | 15 | 11 | $10^5$ | 20 | 1,2,4,7 |
| 49262 | 12 | 11 | $10^5$ | 20 | 1,2,4,7 |
| 40840 | 10 | 10 | $10^5$ | 10 | 4,7 |
| 40827 | 9 | 10 | $10^5$ | 10 | 4,7 |
| 40824 | 11 | 11 | $10^5$ | 10 | 4,7 |
| 36016* | 9 | 3 | $10^5$ | 10 | 2,5,9 |
| 68820 | 28 | 24 | $10^4$ | 10 | 2,4,7 |
| 33266 | 25 | 12 | $10^2, 10^3, 10^4, 10^5$ | 20 | 1,2,4,7 |
| 64660 | 21 | 10 | $10^5$ | 08-Dec | 2,4 |

### D. Antigen Variation on MURINE SARS Gene Expression Analysis GSE49262 and GSE49263

The effects of MA15 on gene expression (GE) were compared to other antigens, including dORF6, NSP16, and Mock antigens, using the datasets GSE49262 and GSE49263 (Figures 4A and B). The Mock samples showed no significant differential GE between 1 DPI (Day Post-Infection) and 7 DPI in both datasets. However, in GSE49262, differential gene expression was observed for VEGF-A (1 DPI versus 2 DPI and 2 DPI versus 4 DPI) and for VEGF-B (1 DPI versus 2 DPI) in the Mock samples. These findings suggest that MA15 elicits a strong immunogenic effect, leading to a significant decrease in GE compared to dORF6 and VEGF-B stimulation. Regarding VEGF-B, only dORF6 (in GSE49262) resulted in a significant decrease in VEGF-B GE over the 4 DPI period. Additionally, dORF6 elicited the v-shaped change in VEGF-B GE mentioned in the previous section. Notably, at the highest dose of MA15, no differential gene expression response was observed for VEGF-B.

In contrast, for both datasets, NSP16 and MA15 induced a decrease in TNF, NFKB1, and VEGF-A GE, with no significant change in VEGF-B GE. Furthermore, compared to dORF6, MA15 stimulation caused a greater decrease in TNF and VEGF-A GE. Similarly, compared to NSP16, NFKB1, and VEGF-A, GE exhibited a more substantial decrease. According to the p-values, the largest decrease in GE magnitude was observed between day 1 and day 4 for MA15 compared to dORF6 and NSP16 for all four genes analyzed using box plot analysis in both datasets.

**Figure 7:**
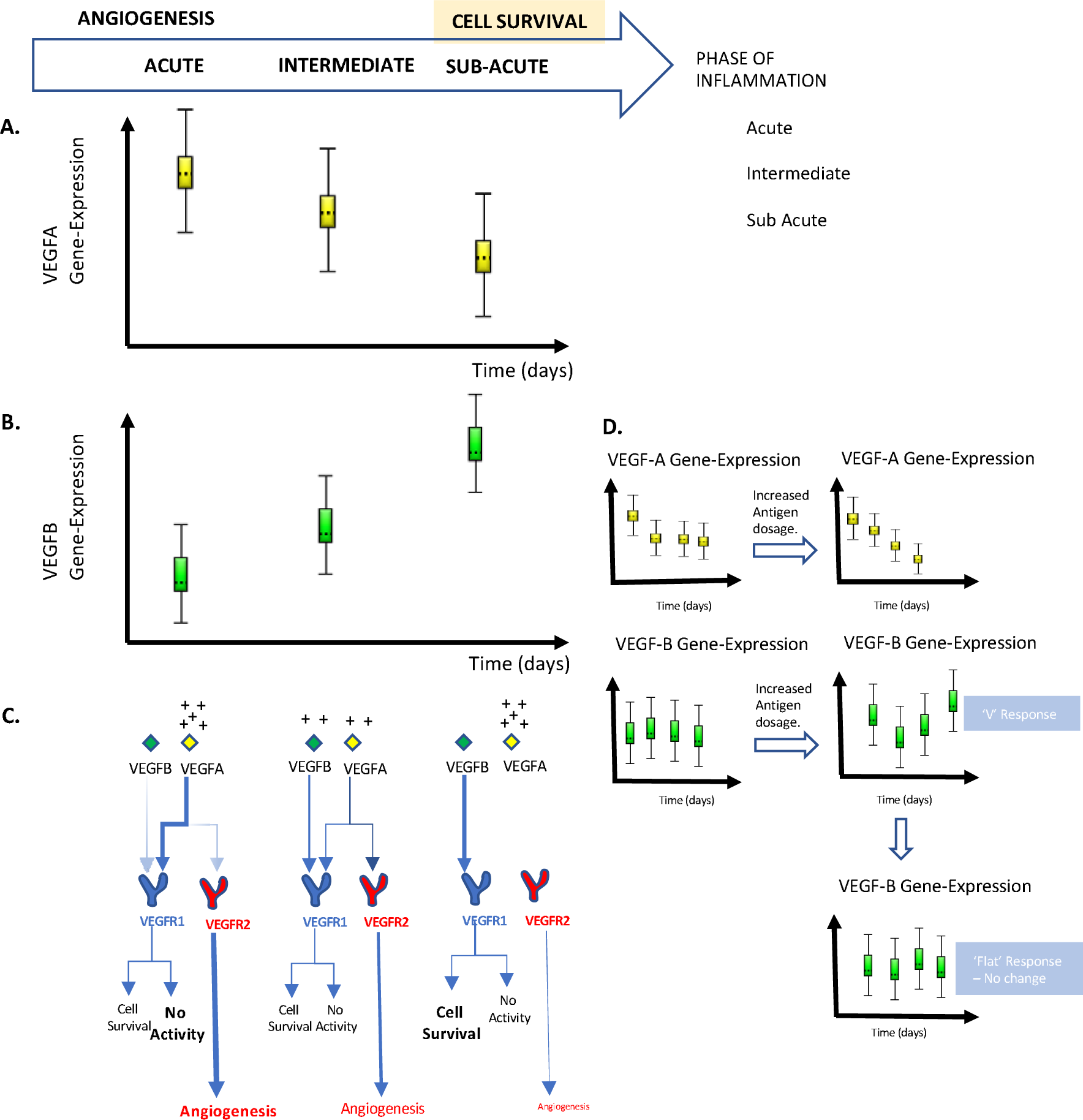
Mechanistic Model of Temporal VEGF-A and VEGF-B changes in Gene Expression, with Pathophysiological Consequences. **Schematic representing a possible model for VEGF-A and VEGF-B interactions Post-SARS-CoV infection in the Pulmonary Murine model.** Temporal changes in VEGF-A and VEGF-B gene expression (GE) are illustrated by way of simplification, depicting trends (Figures 7A and 7B). Three time-associated states are labeled ‘Acute,’ ‘Intermediate,’ and ‘Sub Acute,’ which has implications for whether there is tissue inflammation or whether inflammation has tapered. In the Sub-acute state, we present the idea that molecular pathways are geared toward cell-survival and protection (Figure 7C). It is postulated that there is temporal gene-switching in VEGF-A and VEGF-B GE as the phase of inflammation transitions from one to another. Further, it is suggested that VEGF-A and VEGF-B gene-switching occurs over days. In the acute state, the rise in VEGF-A results in angiogenesis due to VEGFR-2 stimulation. Then in the intermediate state, cell survival and angiogenesis are suggested to be in equipoise. Finally, in the Subacute state, the lower level of VEGF-A results in diminished angiogenesis with enhanced VEGF-B levels resulting in cell survival due to predominate VEGF-B binding to VEGFR-1. Different antigen dosing can affect the temporal gene-expression response, as shown in our paper, which we suggest drives VEGF-A and VEGF-B GE patterns in relation to the degree of inflammation (Figure 7D). We have labeled possible patterns in VEGF-B GE according to our studies on differing doses of MA15 and different antigen types. One possible pattern is the ‘V’ response in VEGF-B GE, and the other is the flat response in VEGF-B GE. The latter response, we suggest, is more likely seen with the higher MA15 (10^5^ PFU) dose and may represent an immune system overwhelmed by the inflammation, being unable to mount a counter-response.

**Figure 8:**
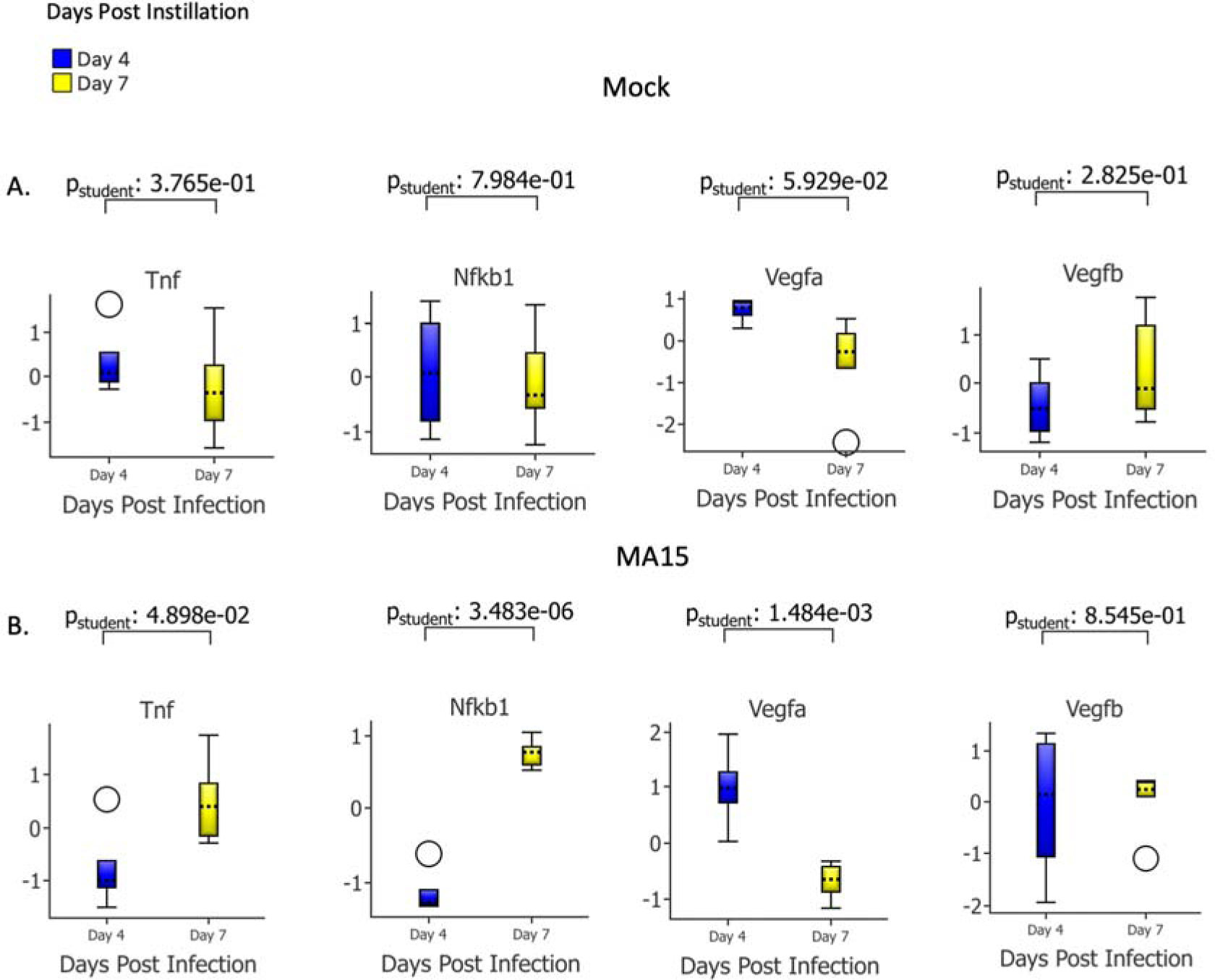
VEGFA, VEGFB Gene Expression, Mock, and WT samples. **GSE40840** Murine pulmonary study showing DGE in the pulmonary bed for MA15 and Mock (10 samples in each group) stimulation in the WT Murine Model. The mock samples showed no differential GE for mock (Figure 8A), whereas for MA15, VEGF-A GE is up-regulated, and down-regulation is noted for VEGF-B GE, TNF, and NFKB1 GE (Figure 8B).

### E. Analysis of changes in TNF, NFKB1, VEGF-A, and VEGF-B in dose and antigen studies (GSE50000)

To further validate the significant changes in TNF gene expression (GE) described earlier, we conducted an additional analysis using an unsupervised approach. We performed UMAP analysis on the dataset GSE50000 to gain insights into the effects of dose and antigen stimulation in the WT Pulmonary Infection Model (Figure 5). The UMAP diagram revealed the presence of three distinct clusters (Figure 5). Two clusters, namely Alpha and Beta, were associated with MA15 samples. Both the MA15 alpha and beta groups exhibited an up-regulation of TNF and a downregulation of VEGF-B. However, the Alpha cluster showed a down-regulation of VEGF-A, while the Beta cluster displayed an up-regulation. Regarding NFKB1, the Beta cluster tended to show downregulation, while the direction of NFKB1 GE for the Alpha cluster was less clear.

Box plots were used to illustrate the GE profiles for the 104 and 105 doses (Figure 9). At the 104 dose, temporal changes in VEGF-B GE were observed, although there were no significant differences between the starting and ending points of the study. However, for the 105 dose, no significant changes in VEGF-B GE were observed, indicating a “flat response” with no notable fluctuations.

### F. PROTEIN-PROTEIN DOCKING

A simulation exercise was performed to investigate the interaction between VEGF receptors and the spike protein of SARS-CoV (Figure 6). The pyDOC platform was utilized to predict the best docking models between VEGF receptors and the spike protein. These models were ranked based on factors such as electrostatics and desolvation energy (Table 3). All predicted models were extracted and plotted to assess the stability of the interactions. Remarkably, the top predicted models exhibited a stable interaction between VEGF receptors and the spike protein of SARS-CoV (Figures 6A-6J).

## DISCUSSION

This study aimed to understand SARS pathogenesis by conducting a temporal transcriptomic analysis focusing on a set of pro-inflammatory genes (TNF, NFKB1, VEGF-A) and VEGF-B, known as the temporal gene set (TGS). Thirteen datasets were systematically procured from the NCBI Geo database, of which seven displayed TGS-related inflammatory changes and were selected for further scrutiny. The relevant studies used MA15, a lethal immunogenic strain of SARS-CoV, was developed for its immunogenicity in the Murine SARS model.

Different murine studies suggest variations in GE timelines. In GSE68820, IL-6 enrichment validated the pro-inflammatory status of the dataset, with TNF and VEGF-A GE diminishing after MA15 exposure, indicating decreasing inflammation. Dataset GSE33266 recorded GE variations at differing MA15 doses. TNF and NFKB1 GE increased early (1-2 DPI) with the 10^3^ and 10^4^ PFU MA15 doses, while VEGF-A declined. Datasets GSE49262 and GSE49263 illustrated declining GE for pro-inflammatory genes, indicating inflammation resolution. Discrepancies between studies may stem from experimental standardization issues and the timing window for analysis. Nevertheless, our research concurs that pro-inflammatory GE should first surge, then decline, while VEGF-A GE downward trends could signify gene-type specific dynamics. Datasets GSE49262 and GSE49263 elucidated differential responses to various antigens, including dORF6, MA15, and NSP16. A notable downward trend in GE was observed for the pro-inflammatory genes (TNF, NFKB1, and VEGF-A) in response to both dORF6 and MA15 antigens, indicating a temporal reduction in inflammation post-stimulation. In comparing the responses from 1-day post-infection (DPI) to 4 DPI, the effect of MA15 stimulation was seen to result in a more significant drop in gene expression for the pro-inflammatory genes under investigation. This effect with MA15 was comparatively more pronounced than the changes observed in response to stimulation with dORF6 and NSP16. Also, MA15 resulted in a flattening of the VEGF-B GE response at the 10^5^ MA15 dose. MA15 is a lethal immunogen in the murine model; one assumes, therefore, that the comparative changes noted in MA15 could be indicative of more severe pro-inflammation. Further, no significant change in VEGF-B response to MA15 could be indicative of a lack of a VEGF-B response. Given that VEGF-B may have a role in cell survival, the lack of response to MA15 suggests an impingement on this function in severe pro-inflammation. Furthermore, an in-depth temporal dose study (GSE50000) revealed that higher doses (10^5^) of MA15 led to a more pronounced decrease in VEGF-A gene expression. Interestingly, VEGF-B showed no change in gene expression at both the 10^4^ and 10^5^ doses. However, at the 104 dose, significant fluctuations in VEGF-B gene expression were observed at different time points, although (as above) the overall gene expression levels at the beginning and end of the study remained not significantly different. This paper thus eluded to interesting patterns of GE in association with pro-inflammation in the WT MA15 Pulmonary model according to the TGS, allowing the document of useful insights. The pattern of VEGF-A and VEGF-B GE temporal association is similar to those described in Sepsis and KD {Rashid, 2022 #4475} (also Sepsis paper in review at the ‘Shock Journal’).

Our research delved into the temporal pattern of VEGF-A and VEGF-B gene expression (GE) post-MA15-induced inflammation. We observed a decrease in VEGF-A GE and an increase in VEGF-B GE over time in GSE68820 following immunogenic stimulation. VEGF-A and VEGF-B, although homologs, displayed distinct temporal patterns across multiple datasets. Dataset GSE33266 showed a declining trend in VEGF-A GE with a concurrent ‘V’ shape trend in VEGF-B GE at the 10^4^ and 10^5^ doses from 1 DPI to 7 DPI. A flat response for VEGF-B GE was noted in some of the datasets at the 105 MA15 dose, indicating a lack of VEGF-B-associated signaling. As VEGF-B production potentially follows acute VEGF-A-induced inflammation, this may have implications for cellular survival. We analyzed transcript profiles of both 104 and 105 MA15 doses in GSE50000. At the 105 doses, VEGF-B GE showed this flat response, while at the 104 doses, VEGF-B GE showed significant fluctuations, although remaining unchanged overall. The study revealed that the antigen dose impacts VEGF-B response and suggests potential biomarkers for inflammation and recovery, highlighting the importance of distinct roles of VEGF-A and VEGF-B.

In this investigation, we employed a pre-selected four-gene set for Differential Gene Expression (DGE) analysis, a decision guided by our previous work. Despite this pre-selection, unsupervised analysis methods were adopted to validate our gene choice and to detect other potentially relevant genes. We utilized one such approach in the GSE68820 dataset, comparing the MA15 infection against a Mock condition. This unsupervised exploration generated a list of differentially expressed genes (DEGs), which further guided us toward related biological pathways. Subsequent enrichment studies, built upon the DEGs, identified connections to inflammation, angiogenesis, and Covid19 processes. The complexity and multidimensionality of the data necessitated the application of Uniform Manifold Approximation and Projection (UMAP) for clustering, enabling differentiation between the Mock and infected samples. This method facilitated labeling subgroups according to the heat-map representation of the genes under study. Distinct patterns began to emerge from the UMAP analysis, most notably the up-regulation of TNF following MA15 infection and the contrasting down-regulation of VEGF-A in certain subgroups. This variation suggests a possible heterogeneity in the rate of VEGF-A decline post-immunogenic stimulation, which could have clinical implications. To further visualize and comprehend the temporal dynamics of gene expression, we applied Temporal Transcriptome Component Analysis (TTCA). This innovative approach reinforced the trends observed in our traditional box plot analyses. For instance, in dataset GSE68820, the TTCA mirrored the observed decline in VEGF-A and the subsequent rise in VEGF-B gene expression. Overall, the combined application of various analytical methods has enriched our understanding of the temporal dynamics of gene expression following MA15-induced inflammation.

This murine transcriptional study corroborates the utility of the MA15 murine pulmonary model for SARS research through the observed enrichment of human SARS-CoV-2 genes (GSE68820). Our investigation suggests an intriguing interplay between the SARS-CoV-2 Spike protein and receptors related to VEGF, hinting at possible pathophysiological processes. For example, the Spike protein may interact with the VEGF-A/neuropilin-1 receptor, resulting in analgesia. Assuming similarities in the interaction of the SARS-CoV MA15 protein receptor with VEGFR1/VEGFR2 and the SARS-CoV-2 receptor, the viral protein may disturb VEGFA/B regulatory feedback mechanisms. This leads us to speculate that the MA15 viral protein might provoke inflammation through VEGF-R2 binding, a hypothesis warranting further research. With an 88% homology between the amino acid sequences of mouse and human VEGF-B, studying VEGF-B may yield important insights.

A key limitation of our study is the assumption that TNF and NFKB1 gene expression (GE) indicate an inflammatory state, guiding our dataset selection. Mock samples, anticipated to denote a non-inflammatory state, serve as a test for this premise. TNF and NFKB1 are known pro-inflammatory genes, so they represent our inflammatory markers. By comparing mock samples to infected ones, we confirmed minimal GE variations (Table 2), supporting Mock’s role as a non-inflammatory state. Aside from one dataset with physiological variation, there were negligible changes in VEGF-A and VEGF-B GE. Thus, the absence of significant GE changes in inflammatory genes in mock samples underscores their role as an unstimulated baseline, highlighting the use of TNF/NFKB1 to classify the mock transcriptome as non-inflammatory. Mock samples are key in refining experimental methodologies, such as quality control and calibration, and we propose a method for this via DGE. Another study limitation relates to the fact that this study does not link physiological parameters with gene expression metrics, an important future research area. Furthermore, while VEGF-A’s role in SARS is well-documented{Chi, 2020 #3336}, with high protein levels linked to severe pulmonary disease in human SARS-CoV-19, our study lacked proteomic data. However, the bioavailability of elevated VEGF-A is difficult to deduce since sFlt-1 acts as a scavenger receptor and neutralizes VEGF-A{Rovas, 2021 #3332}. Therefore, future studies should investigate the connection between GE changes and downstream molecular pathway implications clinically.

**Table 2.** Mock samples and effects on GE for the WT Genotype, stimulated by Mock. In the serial studies, WT mice are given MA15 or mock, and the gene expression from Mock is shown here. Gene expression GE is analyzed for TNF, NFKB1, VEGF-A, and VEGF-B after stimulation and recorded in days. The overall change in GE from the beginning to the end of the study is considered. For dataset GSE68220, a fall in GE in VEGF-A is noted from the start and end of the study are considered (D2 versus D7 p= 4.682e-03).

| GSE | TNF | NFKB1 | VEGF-A | VEGF_B |
| --- | --- | --- | --- | --- |
| 33266 | No change | No change | No change | No change |
| 40840 | No change | No change | No change | No change |
| 49262 | No change | No change | No change | No change |
| 49263 | No change | No change | No change | No change |
| 50000 | No change | No change | No change | No change |
| 68820 | No change | No change | Fall | No change |

This paper introduces the intriguing possibility of modulating VEGF-A and VEGF-B expression after coronavirus infection by adjusting immune stimulant dosages, such as MA15. Both VEGF-A and VEGF-B interact with the VEGFR-1 receptor, but the implications of our temporal findings suggesting a possible inverse relationship between these genes warrant additional research. Pair-wise t-testing may serve as a beneficial complementary approach in this regard. We believe this study’s potential implications are vast. For instance, using gene expression as biomarkers could provide advantages over proteins, especially considering the complex bioavailability of VEGF-A. Moreover, our examination of VEGF-A and VEGF-B changes might have crucial in-vivo modeling and therapeutic implications. For instance, the configuration of VEGF protein-receptor binding can be therapeutically significant, as demonstrated in previous research{Farzaneh Behelgardi, 2018 #4297}{Sadremomtaz, 2020 #4299}. Future therapeutic strategies may focus on the changes and trajectories in VEGF-A and VEGF-B GE. Moreover, Korpela et al. (2022) employed VEGF-B gene therapy on porcine hearts, aiming to promote myocardial benefit{Korpela, 2022 #4305}. Host inflammatory responses undermined the study. However, it does showcase the opportunity to focus on VEGF-B GE. Exploring VEGF-A and VEGF-B gene expression changes in host tissues could yield crucial insights. Our study underscores the importance of considering the cellular interactions and temporal dynamics of VEGF-A/B while targeting angiogenic processes. An important question posed due to GE profiling in this paper is the relationship of the transcript to clinical end-points such as mortality. Future research could focus on the unique dynamics of VEGF-A and VEGF-B and their gene function’s homeostatic feedback system. Moreover, as suggested by our study, the necessity to explore gene expression patterns post-MA15 infection is a crucial avenue for future investigation.

**TABLE 3:**
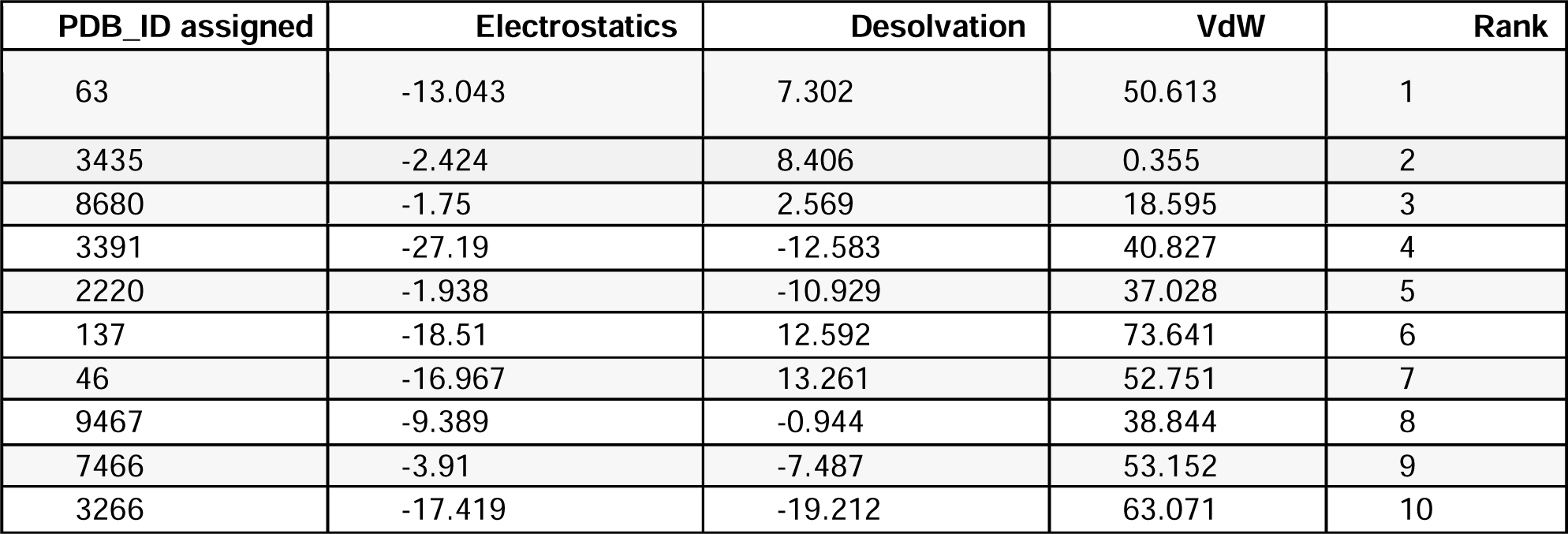
Protein-Protein docking models ranked assigned after docking between VEGF and S Protein of SARS-CoV. A study of VEGF and SARS-CoV protein binding, using Vander wall forces (VdW). The best ten models for VEGF-A and VEGF-B protein modeling with SARS-CoV spike protein was finalized on the basis of best scored by electrostatics and desolvation energy. *In silico* protein-protein interaction signifies that VEGF protein shows stable interaction, which may have pathological consequences.

---

**Supplement Figure 9:**
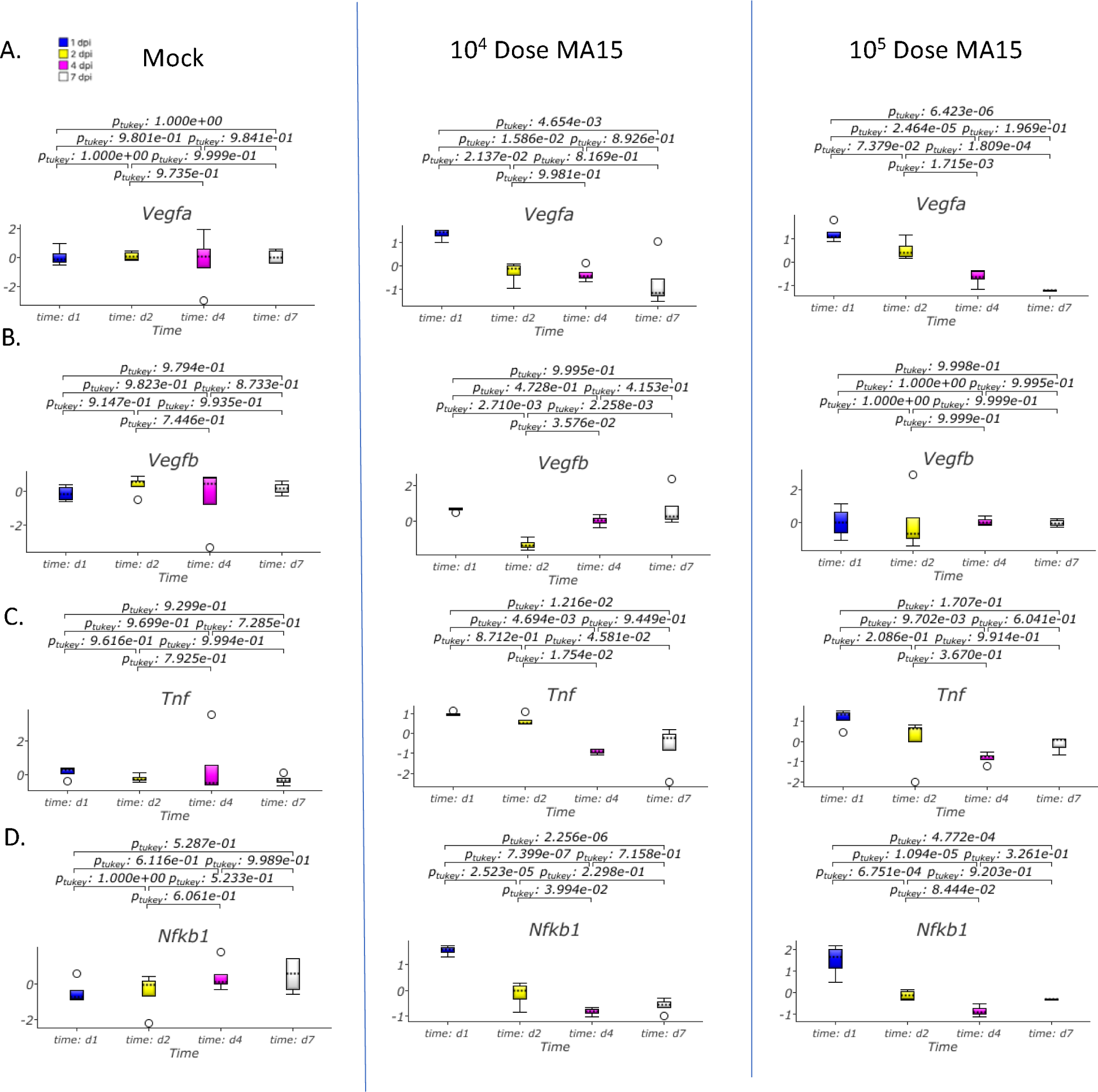
VEGFA, VEGFB Gene Expression, 10^4^ Versus 10^5^ MA15 instillation in the Pulmonary Murine WT Model. Murine MA15 nasal instillation studies are illustrated. Box plots are shown for study datasets GSE50000 (Figure 4A), with gene expression (GE) temporal profiles shown for VEGF-A and VEGFB. Both the 10^4^ and 10^5^ MA15 doses lead to a fall in VEGF-A GE, with the fall being more significant at the 10^5 dose (Figure 5A). There was no change from the mock to the increasing MA15 doses for VEGF-B GE (Day 1 compared to Day 7) (Figure 5B). 16 samples in each group.

